# Components of Mnemonic Metacognition Constitutionally Supported by High Gamma Activity Between the Precuneus and Hippocampus

**DOI:** 10.1101/2024.02.26.581526

**Authors:** Kai Pan, Xinxia Guo, Zhe Zheng, Zhoule Zhu, Hemmings Wu, Junming Zhu, Hongyi Ye, Yudian Cai, Diogo Santos-Pata, Jianmin Zhang, Sze Chai Kwok, Hongjie Jiang

**Author notes:** Correspondence: Hongjie Jiang, Department of Neurosurgery, Second Affiliated Hospital, School of Medicine, Zhejiang University, 88 Jiefang Road, Hangzhou, Zhejiang 310009, China., Sze Chai Kwok, Duke Kunshan University No. 8 Duke Avenue, Kunshan, Jiangsu, 215316 China., Jianmin Zhang, Department of Neurosurgery, Second Affiliated Hospital, School of Medicine, Zhejiang University, 88 Jiefang Road, Hangzhou, Zhejiang 310009, China. Kai Pan and Xinxia Guo authors contributed equally to this work.

## Abstract

Metacognition, the ability to introspectively monitor one’s own performance, remains an area with poorly understood neural mechanisms, particularly regarding the intercommunication among various brain regions. Recent studies have identified the precuneus as a key region in mnemonic metacognitive tasks, while the hippocampus is recognized for its role in memory function. This study aimed to investigate the role of the precuneus and hippocampus in mnemonic metacognition by utilizing intracranial electrode recordings from patients with intractable epilepsy to analyze the correlational dynamics between these regions. Our findings revealed that high gamma activity serve as a key feature in both the precuneus and hippocampus for confidence judgment. Based on signal detection theory (SDT), we discovered that high gamma activity in the precuneus is significantly linked to type 2 sensitivity (meta-d’), while hippocampal high gamma activity was primarily linked to type 1 sensitivity (d’). Additionally, the correlation between high gamma activity in the hippocampus and precuneus was exclusively related to type 2 sensitivity (meta-d’). Temporal analysis indicated that the hippocampus is initially engaged for the memory process, followed by its joint engagement together with the precuneus for confidence generation. These findings elucidate the distinct electrophysiological roles of the hippocampus and precuneus in mnemonic metacognition, providing deeper insights into the neural mechanisms underlying this cognitive process.

## INTRODUCTION

Metacognition, defined as the ability to introspectively monitor and regulate cognitive processes, involves reevaluating cognition and objectively assessing confidence in decision-making (Flavell, 1979; Fleming et al., 2024; Miyoshi et al., 2024). Numerous studies have elucidated the neural underpinnings of various domains of metacognition, particularly in relation to perception and memory. These investigations have highlighted that the prefrontal cortex’s association with perception-related metacognition, and that the precuneus is more intricately linked to memory-related metacognition (Fleming et al., 2010; McCurdy et al., 2013; Baird et al., 2013). Despite advances in identifying the critical roles of the prefrontal (Hebart et al., 2016; Shekhar and Rahnev, 2018; Cai et al., 2022) and parietal cortices (Morales et al., 2018; Pereira et al., 2021) in metacognitive processes, the interregional communication underlying the computation for metacognition remain elusive.

The precuneus has been implicated not only in metacognitive processes related to memory but also in broader memory functions. Studies have demonstrated its involvement in maintaining object-location within working memory (Bicanski & Burgess, 2018), facilitating the retrieval of autobiographical memories (Hebscher et al., 2020), and reconstructing contextual information to aid temporal order memory tasks (Foudil et al., 2020). Given the precuneus’s integral role in memory-related cognitive processes and its anatomical connection with the hippocampus (Tanglay et al., 2022), we hypothesized the existence of a neural circuit linking the precuneus and the hippocampus. Empirical evidence supports this hypothesis, documenting hippocampal-precuneus connections that can predict memory retrieval confidence levels (Ren et al., 2018) and highlighting the importance of hippocampal-posterior parietal cortex information flow in episodic memory tasks (Das and Menon, 2023). The pathway between the hippocampus and the precuneus has also been implicated in reasoning decision processes (Pudhiyidath et al., 2022) and in self- and goal-oriented processing (Zheng et al., 2021). The involvement of hippocampal-parietal circuits in these related processes suggests that communication between the hippocampus and precuneus is also crucial for memory-related metacognition.

Previous research, which has predominantly utilized magnetic resonance imaging, animal neurophysiology, or cognitive alterations under pathological states, has yet to ascertain alterations in neural electrical activity within the human brain during mnemonic metacognitive processes. Our study addressed this lacuna by analyzing intracranial electroencephalogram signals from patients with intractable epilepsy, uncovering the correlation and potential neural pathways linking the precuneus and hippocampus within mnemonic metacognition. To gain a better understanding of metacognition and the roles of the precuneus and hippocampus within this performance, our study employs methodologies based on the signal detection theory (SDT) framework (Maniscalco and Lau, 2012). By integrating behavioral and electrophysiological signal analyses, our study elucidated the unique neural mechanisms of metacognition, thereby contributing to a deeper understanding of this cognitive process.

## RESULT

A total of eleven patients with intractable epilepsy underwent stereoelectroencephalography (sEEG) surgery, during which their brain electrical signals were recorded during metacognitive behavioral tasks. The demographic and clinical characteristics of eleven subjects are provided in Supplementary Table S1. Postoperative metacognitive behavioral testing was conducted in the ward, where patients were first shown a video and subsequently asked to determine the temporal order of two displayed pictures. For each trial, after the memory judgment, they were required to make a confidence judgment (Fig. 1a; see Methods section for details). Schematic diagram of electrodes was illustrated in Fig.1b. To ensure accurate electrode placement, we calculated the Montreal Neurological Institute (MNI) coordinates and visualized them using the AAL3 atlas (Fig. 1c).

**Figure 1.**
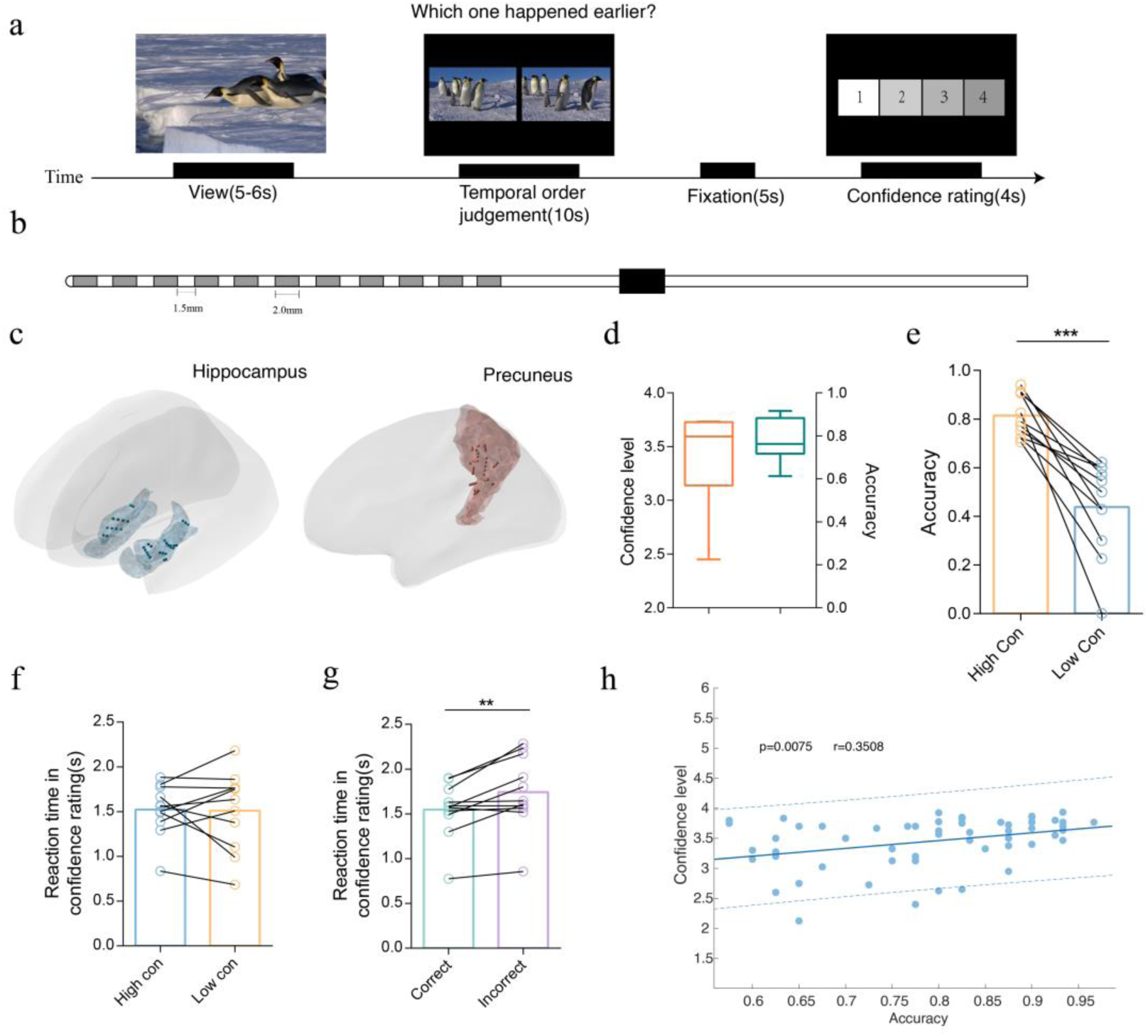
Experimental design, electrode’s location and behavioral details. **(a)** Experimental design. Participants performed a temporal order judgment task, by choosing the image that happened earlier in the video. After this, participants needed to report confidence level from 1-4 points. **(b)** Schematic diagram of stereoelectroencephalography electrodes used in this study. **(c)** A summary of the recording locations from 11 participants. Each colored dot represents the location of a recording site shown on a standard brain template (AAL Atlas). The two colors represent the brain areas. Blue: hippocampus; Red: precuneus. **(d)** Bar plot shows average confidence level (3.40±0.41) and average accuracy (77%±9.5) across participants. **(e)** Accuracy plotted in high confidence and low confidence trials across participants, with a significant difference (paired t-test, p = 0.0078). **(f)** Reaction time in confidence rating stage of high confidence trials compared with low confidence trials across participants (paired t-test, p > 0.05). **(g)** Reaction time in confidence rating stage of correct trials is significantly shorter than incorrect trials (paired t-test, p = 0.0058). **(h)** Correlation between the confidence level and accuracy across sessions collapsed over participants (r = 0.3508, p = 0.0075). Each dot denotes a session.

### Behavioral results

We calculated the mean levels of confidence (3.4148 ± 0.1206 mean ± SEM) and accuracy (0.7738 ± 0.0288 mean ± SEM) for these eleven participants during the behavioral experiment (Fig. 1d). After classifying the trials according to the level of confidence, we found that trials with high confidence had significantly higher accuracy compared to trials with low confidence (Fig. 1e, p < 0.001, paired t-test). Reaction times during the confidence stage were significantly shorter in correct trials than in incorrect trials (Fig. 1g, p < 0.01, paired t-test), although no significant difference was observed between high confidence and low confidence trials (Fig. 1f, p > 0.05, paired t-test). There was a significant positive correlation between accuracy and confidence level across sessions (Fig. 1h, r = 0.3508, p = 0.0075, Pearson correlation).

#### Precuneus and hippocampus involvement in metacognition through low gamma and high gamma frequency band

We first determined the frequency bands in the hippocampus and precuneus that are most critical for and involved in metacognitive processes. We conducted a time-frequency analysis, comparing the fixation stage with the confidence stage to calculate t-values for these regions (Fig. 2a, 2e). The results revealed significant increased activation in the precuneus and the hippocampus in both low gamma γ (35-70 Hz) and high gamma Hγ (70-150 Hz) frequency bands. To assess the high frequency band impact of the precuneus and hippocampus on confidence ratings, we employed generalized linear models (GLMs) to regress the γ and H γ bands against the confidence level. We identified significant differences in the -1.0 to -0.5 s interval for both the hippocampus and precuneus (Fig. 2b, 2f, cluster-based permutation, p < 0.05). Additionally, to verify the contribution of each frequency band to confidence, we compared GLMs that included only high-gamma and gamma bands with all possible GLMs that included these high-frequency bands plus any combination of low-frequency bands. Bayesian model selection (see Methods) indicated that the high-frequency bands (both low and high gamma) GLM provided the best account of subjective value (exceedance probability (Xp) = 1). Regression coefficients for each band showed significant differences for high gamma compared to other bands (Fig. 2c, 2g, one-sample, two-sided t-test, p < 0.05). To further explore the influence of other task-related factors on high-frequency activity, we performed a regression analysis on the γ and Hγ bands. The results indicated that, compared to other factors, confidence level and confidence reaction time had a more pronounced impact. (Fig. 2d, 2h, one-sample, two-sided t-test, p < 0.05). These results suggest that both the hippocampus and precuneus contribute to confidence rating within the high gamma band, displaying similar time windows of activity.

**Figure 2.**
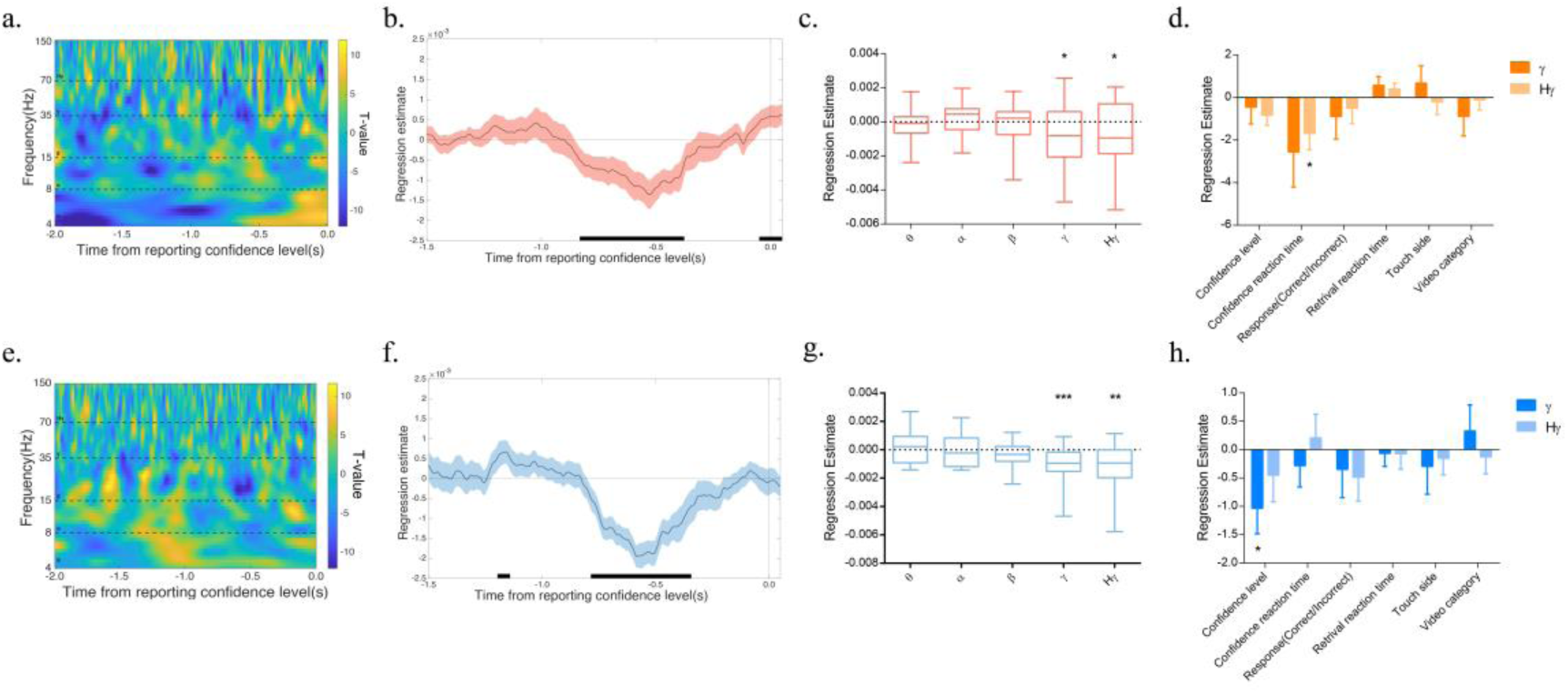
Gamma activity in the precuneus and hippocampus associated with confidence. **(a)** Time-frequency analysis (T-value map) during the confidence rating stage, compared with the fixation stage, averaged across all electrodes in the precuneus. Horizontal dashed lines indicate boundaries between frequency bands that are included in the analysis in (c). **(b)** Time course of regression estimates for the gamma and high-gamma frequency bands in the precuneus. The solid line (shaded areas) represents the SEM across the precuneual electrode sites (n=31). The horizontal black rectangle on the x-axis represents significant time points (cluster-based permutation, p < 0.05). **(c)** Regression estimates of power against confidence rating in the precuneus, averaged over the - 1.0 to -0.25s window, for each frequency band defined in (a). Asterisks indicate significance of regression estimates (one-sample, two-sided t-test, p < 0.05). **(d)** Regression estimates of related factors against gamma and high gamma in the precuneus, averaged over the -1.0 to -0.25s window. Asterisks indicate significance of regression estimates (one-sample, two-sided t-test). **(e)** Time- frequency analysis (T-value map) during the confidence rating stage, compared with the fixation stage, averaged across all electrodes in the hippocampus. Horizontal dashed lines indicate boundaries between frequency bands that are included in the analysis in (e). **(f)** Time course of regression estimates for the gamma and high-gamma frequency bands in the hippocampus. The solid line (shaded areas) represents the SEM across the hippocampal electrode sites (n=24). The horizontal black rectangle on the x-axis represents significant time points (cluster-based permutation, p < 0.05). **(g)** Regression estimates of power against confidence rating in the hippocampus, averaged over the -1.0 to -0.25s window, for each frequency band defined in (e). Asterisks indicate significance of regression estimates (one-sample, two-sided t-test, γ: p < 0.01, Hγ: p < 0.001). **(h)** Regression estimates of related factors against gamma and high gamma in the hippocampus, averaged over the -1.0 to -0.25s window. Asterisks indicate significance of regression estimates (one-sample, two-sided t-test, p < 0.05).

#### High gamma activity reflects distinct cognitive processes in the precuneus and hippocampus

Given the involvement of high gamma band activity in the confidence rating process, we investigated the cognitive processes represented by high gamma activity in the precuneus and hippocampus. Metacognition is often measured by calculating d’ and meta-d’ based on signal detection theory (SDT). To explore this, we computed an accuracy index with the neural data (Hγ _Correct_ – Hγ _Incorrect_) which corresponds to the behavioral index d’. We also calculated an interaction index of confidence and accuracy using the neural data [(Hγ _High confidence_ – Hγ _Low confidence_) - (Hγ _Correct_ – Hγ _Incorrect_)] to correspond to the behavioral index meta-d’, which refers to one’s efficacy with which confidence ratings discriminate between correct and incorrect judgments.

Using a 0.1s window, we assessed changes in power spectral density of high gamma activity during the confidence rating stage, standardizing the data using z-scores. In the precuneus, high gamma activity began to change between approximately -0.3 s and 0 s, for either comparing high vs. low confidence or comparing high vs. low accuracy trials (Fig. 3a, left and right panels). In contrast, in the hippocampus, changes occurred between -0.8 s and -0.5 s, also showing consistency for both confidence and accuracy (Fig. 4a, left and right panels).

**Figure 3.**
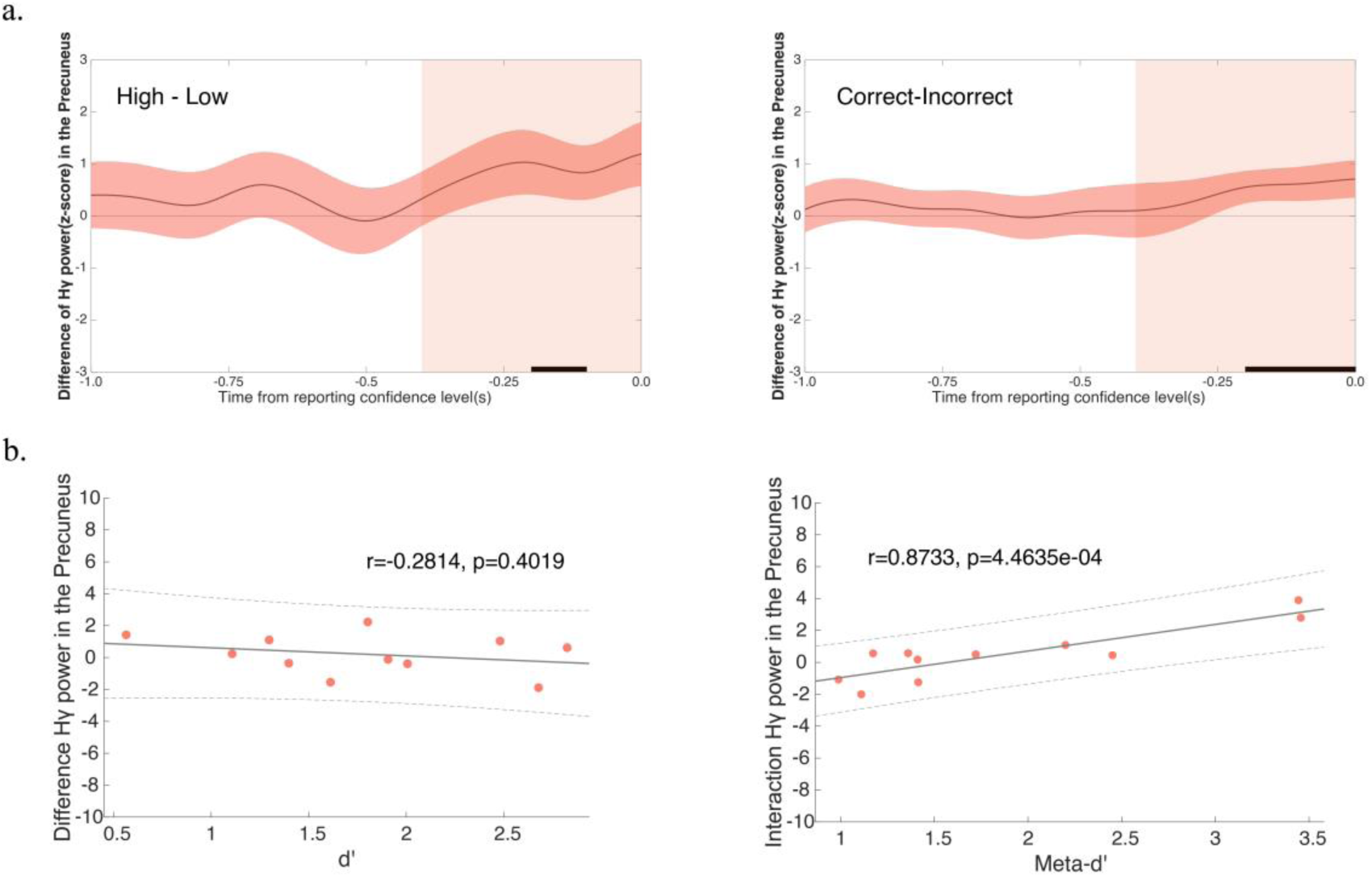
High gamma power in the precuneus is associated with meta-d’ but not with d’. **(a)** Left panel: The time course of the difference in high gamma power (z-score) of the precuneus between high confidence trials and low confidence trials. Right panel: The time course of the difference in high gamma power (z-score) of the precuneus between correct trials and incorrect trials. The solid line represents the mean with SEM (shaded areas) across electrode sites (n=31). The horizontal black rectangle on the x-axis represents significant time points (cluster-based permutation, p < 0.05). The vertical red shaded areas denote the time window of interest used in **(b)**. **(b)** The left panel illustrates the correlation between the neural accuracy index (Hγ _Correct_ – Hγ _Incorrect_) and behavioral type 1 sensitivity (d’) (Pearson correlation, r = -0.2814, p = 0.4019), whereas the right figure depicts the correlation between the neural interaction index [(Hγ _High confidence_ – Hγ _Low confidence_) - (Hγ _Correct_ – Hγ _Incorrect_)] and behavioral type 2 sensitivity (meta-d’) (Pearson correlation, r = 0.8733, p < 0.001).

**Figure 4.**
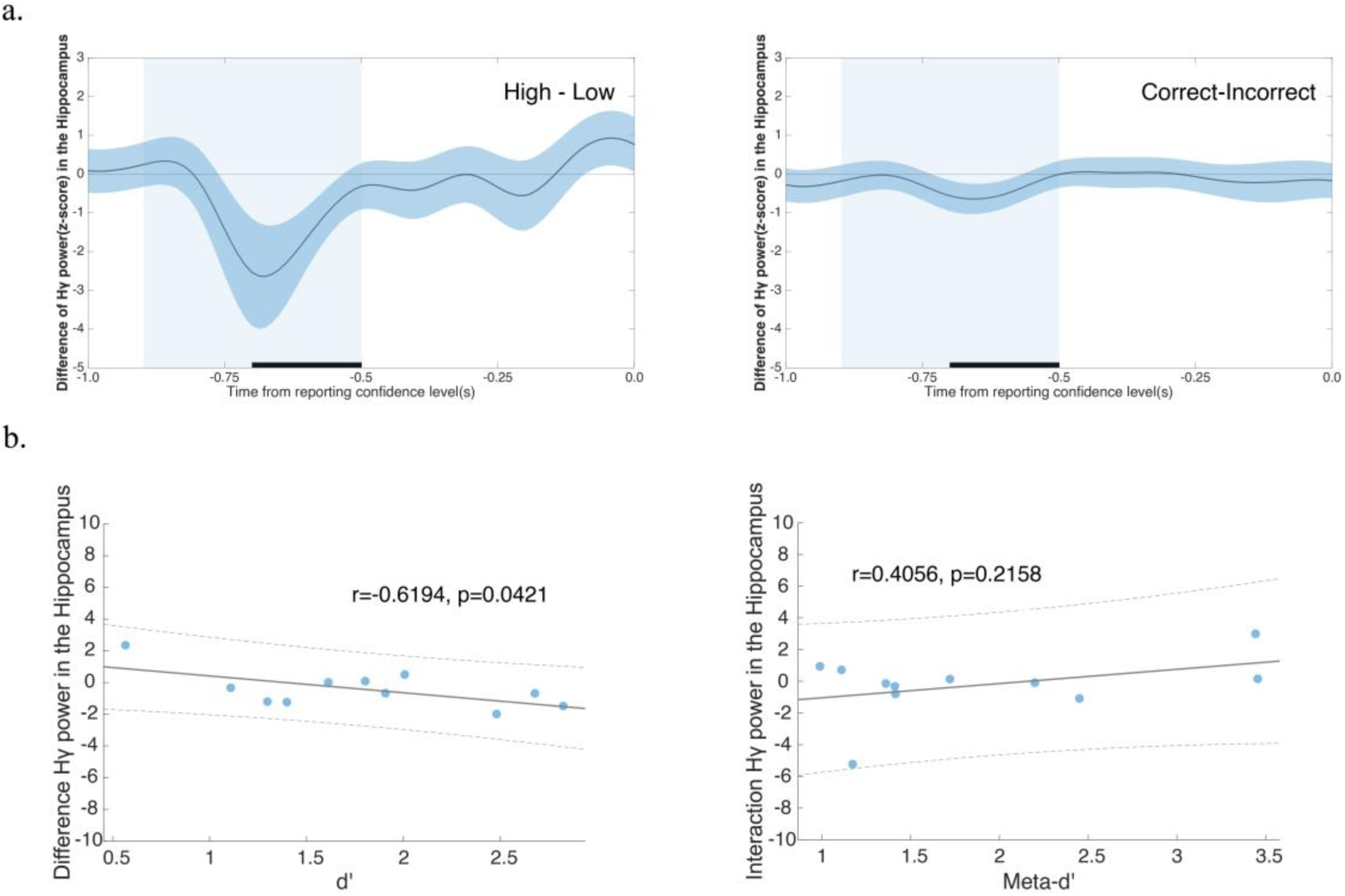
High gamma power in the hippocampus is associated with d’ but not with meta-d’. **(a)** Left panel: The time course of the difference in high gamma power (z-score) of the hippocampus between high confidence trials and low confidence trials. Right panel: The time course of the difference in high gamma power (z-score) of the hippocampus between correct trials and incorrect trials. The solid line represents the mean, with SEM (shaded areas) across electrode sites (n=24). The horizontal black rectangle on the x-axis represents significant time points (cluster-based permutation, p < 0.05). The vertical shaded areas highlight the time window of interest used in **(b)**. **(b)** The left panel illustrates the correlation between the neural accuracy index (Hγ _Correct_ – Hγ _Incorrect_) and behavioral type 1 sensitivity (d’) (Pearson correlation, r = -0.6194, p = 0.0421), whereas the right panel depicts the correlation between the neural interaction index [(Hγ _High confidence_ – Hγ _Low confidence_) - (Hγ _Correct_ – Hγ _Incorrect_)] and behavioral type 2 sensitivity (meta-d’) (Pearson correlation, r = -0.4056, p = 0.2158).

We then averaged the accuracy index and interaction index of high gamma within these two windows across all subjects and conducted Pearson correlation analyses with d’ and meta-d’, respectively. Pearson correlation analyses revealed that high gamma activity in the precuneus was significantly correlated with meta-d’, but not with d’ (Fig. 3b). Conversely, high gamma activity in the hippocampus was significantly correlated with d’, but not with meta-d’ (Fig. 4b). Further analysis showed no significant correlation between precuneus high gamma activity during the hippocampus’s significant time window and d’ or meta-d’ (Supplementary Fig. S1a). However, hippocampal high gamma activity during the precuneus’s significant time window showed a trend towards correlation with meta-d’ (Supplementary Fig. S1b). These findings suggest that the hippocampus and precuneus both play a role in confidence judgment, with the precuneus being more involved in metacognitive processes (reflected by meta-d’) and the hippocampus more engaged in memory processes (reflected by d’). The different time windows emphasize their distinct contributions. Given the observed trend of correlation between high gamma activity in the hippocampus and meta-d’ within the same time window, our next step is to investigate whether the activities between the hippocampus and precuneus might be correlated or synchronized differently for these distinct cognitive processes.

#### Correlation between high gamma activity in the precuneus and hippocampus related with metacognition

Having established the relationship between the high gamma activity in the precuneus and metacognition, we next examined how hippocampal gamma activity might correlate with that of the precuneus. We applied a correlation analysis method using a 0.02 s window and 0.01s steps to compute Pearson correlation coefficient between the hippocampus and precuneus signals. Amplitude segments with significant correlations lasting over 0.04 s were counted (Fig. 5a). Additionally, segments lasting over 0.06 s were calculated, showing results consistent with those obtained with using 0.04 s segments (Supplementary Fig. S2). We ran correlation analyses for each pair of electrode sites (82 pairs in total) from the 11 subjects. Overall, we found that 60 out of these 82 electrode pairs between the precuneus and hippocampus showed significantly higher correlation values compared to shuffled data. Taking one example participant, the original data exhibited a higher probability density in greater counts than the shuffled data (Fig. 5b, p < 0.001, permutation test, n = 1000). We then ran this analysis on each participant individually and found the same significant differences (Supplementary Fig. S3). The correlation count for the original data was significantly higher than that for the shuffled data (Fig. 5c, Wilcoxon matched pairs signed-rank test, p < 0.0001). This elevated correlation count indicates a stronger relationship between high gamma activity in the hippocampus and precuneus. Moreover, we applied this index to examine its relationship with the metacognitive indices. When categorized by confidence level, high confidence trials had a significantly higher correlation count than low confidence trials (Fig. 5d, Wilcoxon matched-pairs signed-rank test, p < 0.05). Incorrect trials exhibited a significantly higher correlation count than correct trials (Fig. 5e, Wilcoxon matched-pairs signed-rank test, p < 0.0001). In alignment with the power analysis, we also calculated the interaction index in correlation count [(Count _High confidence_ – Count _Low confidence_) - (Count _Correct_ – Count _Incorrect_)], which was significantly greater than zero (Fig. 5g, one-sample two-sided t-test, p < 0.0001). For completeness, we ran Pearson correlation analyses and revealed that the interaction index showed a trend towards correlation with meta-d’ (Supplementary Fig. S4b) but not between the accuracy index and d’ (Fig. S4a).

**Figure 5.**
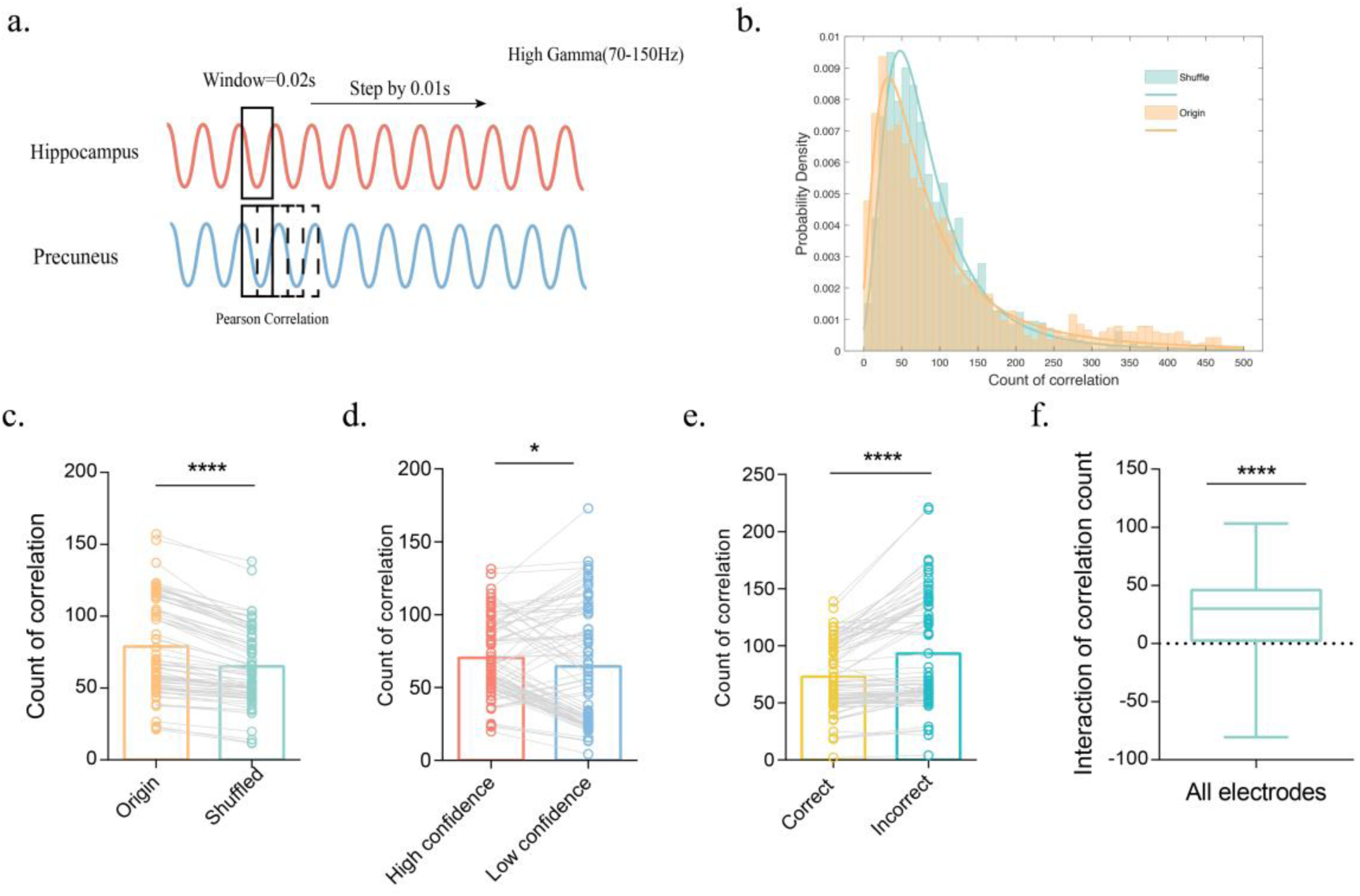
Correlation analysis of high gamma activity between the precuneus and the hippocampus and its relationship with metacognitive indices. **(a)** Schematic diagram of correlation analysis procedure. We defined a 0.02s length window, step by 0.01s, and calculated Person correlation in each window. Segments in which 4 or more consecutive windows (0.04s, at least 2 cycles of high gamma) showing significant correlation are considered high correlation. We calculated the count of highly correlated occurrences between the hippocampus and the precuneus. **(b)** The probability density of one representative participant comparing shuffled and original data is presented (p < 0.001, permutation test, n=1000), see also Figure S2. **(c)** Count of correlation of original and shuffled data (Wilcoxon matched pairs signed rank test, p < 0.0001) taking 82 electrode pairs electrodes from all participants into account. **(d)** Count correlation in high confidence trials compared with low confidence trials (Wilcoxon matched pairs signed rank test, p < 0.05). **(e)** Count correlation in correct trials compared with incorrect trials (Wilcoxon matched pairs signed rank test, p < 0.0001). **(f)** Neural Interaction index [(Count _High confidence_ – Count _Low confidence_) - (Count _Correct_ – Count _Incorrect_)] of high correlation count across all electrodes against zero (one-sample t-test, p < 0.0001). SEM is across all electrode pairs.

Given the strong correlation identified between the precuneus and hippocampus during the confidence rating stage, we then examined the temporal dynamics of this relationship. We calculated the Pearson correlation coefficient between the amplitudes of the hippocampus and the precuneus using a 0.02s window, averaging it according to the 0.1s window used in power analysis. We called this correlation coefficient as H-P correlation. The H-P correlations were significant at approximately -0.3 s and -0.9 s (Fig. 6a), aligning with previously identified changes in the power analysis (Fig. 3a, 4a). Additionally, we conducted Pearson correlation analyses between the H-P correlation and either d’ or meta-d’. The H-P correlation coefficients at approximately - 0.9s showed no significant correlation with either meta-d’ or d’ (Fig. 6b). In contrast, the H-P correlation coefficients around -0.3s exhibited a significant correlation with meta-d’ but were not with d’ (Fig. 6c). These results demonstrated that the precuneus and hippocampus might jointly participate in the process of confidence rating, with their interaction being specifically related to type 2 performance rather than type 1 performance, highlighting the significance of their connection in metacognitive processes.

**Figure 6.**
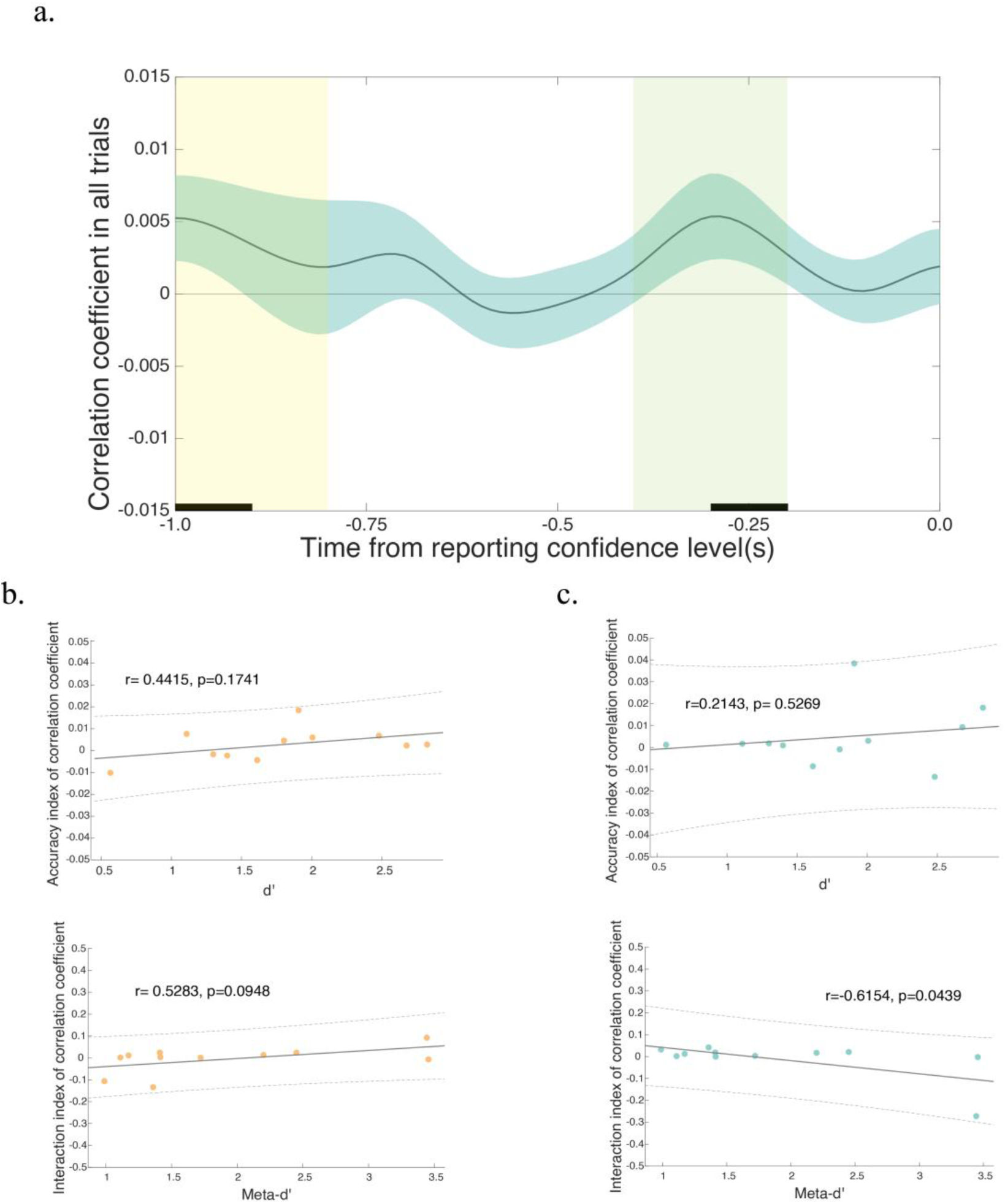
The time course of correlation coefficient of high gamma between the precuneus and the hippocampus towards confidence level reporting. **(a)** The time course of correlation coefficient before confidence rating in all trials. The solid line represents the mean with SEM (shaded areas) across all subjects (n=11). The horizontal black rectangle on the x-axis represents significant time points (cluster-based permutation, p < 0.05). The yellow and green vertical shaded areas represent the time window of interest used in **(b)** and **(c)** respectively. **(b)** The upper panel illustrates the correlation between behavioral type 1 sensitivity (d’) and neural accuracy index (H-P _Correct_ – H-P _Incorrect_) of H-P correlation coefficient (Pearson correlation, r = 0.4415, p = 0.1741), whereas the lower panel depicts the correlation between behavioral type 2 sensitivity (meta-d’) and neural interaction index [(H-P _High confidence_ – H-P _Low confidence_) - (H-P _Correct_ – H-P _Incorrect_)] of H-P correlation coefficient (Pearson correlation, r = 0.5283, p = 0.0948). **(c)** The upper panel illustrates the correlation between behavioral type 1 sensitivity (d’) and neural accuracy index (H-P _Correct_ – H- P _Incorrect_) of correlation coefficient (Pearson correlation, r = 0.2143, p = 0.5269), whereas the lower panel depicts the correlation between behavioral type 2 sensitivity (meta-d’) and neural interaction index [(H-P _High confidence_ – H-P _Low confidence_) - (H-P _Correct_ – H-P _Incorrect_)] of H-P correlation coefficient (Pearson correlation, r = -0.6154, p = 0.0439).

## Discussion

In this study, we employed intracranial electrode recordings to investigate changes in brain electrical activity during mnemonic metacognitive processes. Specifically, we analyzed first-order and second-order performance in metacognition by examining the distinct characteristics and correlations of specific frequency bands in the precuneus and hippocampus. Our findings underscore the critical role of the temporal-to-parietal pathway in the genesis of metacognition, emphasizing the significance of neural circuits over isolated brain regions in metacognitive processes.

Our study is the first to identify the involvement of high gamma band activity in both the precuneus and hippocampus during the confidence judgment stage. The role of the precuneus in metacognition, including mnemonic metacognition, has been extensively documented in previous research (McCurdy et al., 2013; Baird et al., 2013; Ye et al., 2018; Ye et al., 2019), as well as in non-memory domains (Fleming et al., 2010; Heereman et al., 2015; Morales et al., 2018). The literature suggests that high gamma activity in the precuneus is associated with memory retrieval processes (Hebscher et al., 2019), whereas our findings provide evidence that the precuneus contributes to metacognitive processes via high-frequency bands (Manippa et al., 2024). This may explain the cognitive improvements observed following high-frequency stimulation of the precuneus in dementia patients (Morris and Hannesdottir, 2004; Mograbi et al., 2012; Koch et al., 2022).

Metacognition, as a complex and advanced cognitive function, should not be anatomically attributed to a single brain region. Indeed, like in memory, mnemonic processes are multifaceted and can be neurally dissociated into precision, vividness, and accuracy (Richter et al., 2016). Beyond that, our ability to monitor some of these memory-related processes is also segregated neurally (Zou and Kwok, 2022). Previous studies have suggested the role of functional connectivity between the precuneus and the hippocampus in metacognitive processes. The two regions are linked by the superior longitudinal fasciculus (SLF), emphasizing the relevance of both white and grey matter pathways (Zheng et al., 2021). Building on that, this study focuses on the functional connection between the precuneus and the hippocampus. Several studies have identified significant interactions between the hippocampus and precuneus in various cognitive functions, including memory (Ren et al., 2018), reasoning and decision-making (Pudhiyidath et al., 2023), and self- and goal-oriented processing (Zheng et al., 2021). Regarding the neural mechanisms underlying these interactions, phase-amplitude coupling has been observed between the medial temporal lobe and precuneus during memory retrieval (Hebscher et al., 2019). Our study supports the involvement of both the hippocampus and the precuneus in metacognitive processes at high-frequency bands, highlighting potential differences in the neural mechanisms underlying memory retrieval and metacognition. As a control, we conducted a phase-amplitude coupling (PAC) analysis between the hippocampus and precuneus (Voytek et al., 2009; Jensen et al., 2016), which showed no significant difference in coupling between the retrieval and confidence stages (Supplementary Fig. S5). Although this finding provides no direct evidence for confidence-related PAC, we propose that this PAC phenomenon likely originates from memory retrieval processes (Richter et al., 2016).

Pereira et al. (2021) reported a strong correlation between evidence accumulation and meta-perception in the posterior parietal cortex by single-neuron activity of a microelectrode. However, the limitations of electrocorticography (ECoG) in their study left unresolved whether deep brain structures, such as the hippocampus, are implicated during metacognition. There is evidence suggesting the neural circuitry linking the hippocampus to the cortex is important for metacognition. Some work reported that the structural integrity of the fornix, as the output tract of the hippocampus, is related to metacognition driven cognitive offloading (Zheng et al., 2024). This suggests that the fornix is not only important for mnemonic functions (Kwok et al., 2015; Kwok & Buckley, 2010) but also provides a pathway to deliver memory related signals for the frontal-parietal cortice (Boldt & Gilbert, 2022; Kwok et al., 2019) s to generate confidence and metacognitive computation. These are in line with reports that the hippocampus plays a role in memory-related metacognition (Allen et al., 2017; Miyamoto et al., 2017, 2018). This possibility is corroborated with findings showing resting-state functional connectivity (rs-fcMRI) between the precuneus and hippocampus is positively correlated with metamemory ability (Ye et al., 2019). This is in line with anatomical evidence for the interaction between the two regions as they are connected by the fiber bundle between the precuneus to the parahippocampal gyrus (Tanglay et al. 2022). Our findings thus help characterize the interaction via these cortical-subcortical regions in metacognitive processes in a memory task.

In the hippocampus, many studies have demonstrated the role of high gamma bands in memory-related cognitive activities (Burke et al., 2014; Staresina et al., 2016; Mart, 2021). In addition to their involvement in memory processes, memory-selective neurons in the hippocampus exhibit firing patterns associated with confidence judgments (Rutishauser et al., 2015). Gray matter myelination in the hippocampus has been linked to individuals’ metacognitive abilities (Allen et al., 2017). Our findings suggest that high gamma activity in the hippocampus also plays a role in metacognitive processes. Given the joint involvement of the precuneus and hippocampus in confidence reporting, the correlation between high gamma activity in these two brain regions is particularly noteworthy. Indeed, our study confirms a significant correlation between high gamma activity in the precuneus and the hippocampus.

Importantly, our results showed first-order and second-order performance in metacognition could be dissociated through a high gamma band communication between the hippocampus and precuneus. In memory tasks, first-order performance refers to the memory performance. We identified a significant correlation between first-order performance and high gamma band power in the hippocampus, with this activity occurring approximately 0.8 to 0.5 s before the confidence reporting. Additionally, elevated gamma band activity in the hippocampus during confidence reporting was observed for trials with incorrect and high confidence responses. The high gamma band activity was negatively correlated with type 1 sensitivity (d’) across participants, suggesting that the hippocampus may generate an "error signal" during first-order performance, consistent with previous findings (Haque et al., 2020; Desender et al., 2021). Regarding confidence, or second-order performance, we found that it to be primarily correlated with high gamma power in the precuneus, occurring approximately 0.3 to 0 s before the confidence rating. A similar time window, spanning 0.2 to 0.5 s after stimulus onset, had been observed in meta-perception (Xue et al., 2023). Importantly, the correlation between the precuneus and hippocampus was significantly associated with second-order performance but not with first-order performance. These findings confirm that the hippocampus and precuneus cross talk to each other and jointly contribute to metacognition. While we established the synergistic role of the precuneus and hippocampus in metacognitive processes, we furthermore observed some functional differences between the two regions. Neither the precuneus alone nor its interaction with the hippocampus was associated with first-order performance; only the hippocampus showed involvement in these first-order processes. We also observed significant correlation around the first-order performance period of hippocampus, which may reflect functional connectivity between the precuneus and hippocampus during working memory processes (Ren et al., 2018; Hebscher et al., 2019). These findings underscore the distinct roles of the hippocampus and precuneus in metacognition process: the hippocampus contributes to both memory and metacognitive process, while the precuneus appears specifically engaged in metacognitive processes. These results also highlight a temporal distinction between first-order performance (d’) and metacognitive performance (meta-d’). It would be important to further verify whether this temporal distinction is domain-specific or generalizable across different types of metacognition in future research.

In conclusion, our study demonstrates that the precuneus and hippocampus are both implicated in mnemonic metacognition. Through high gamma band activity communication, the precuneus and hippocampus together contribute to second-order performance, whereas first-order performance is primarily mediated by the hippocampus alone. Our findings provide electrophysiological evidence of a parietal-hippocampal circuit (see also the role of fornix in Zheng et al., 2024) (see also the role of fornix in Zheng et al., 2024) underpinning mnemonic metacognition in humans.

## METHODS AND MATERIALS

### Participants

Eleven patients diagnosed with intractable epilepsy underwent implantation of subdural intracranial electrodes for epilepsy treatment at The Second Affiliated Hospital, Zhejiang University School of Medicine (SAHZU). The patients’ ages ranged from 12 to 39 (mean=24.91 years, SD=9.63). All patients demonstrated a good understanding of the experimental procedure and had at least 1 electrode recording signals in each of the target brain regions (hippocampus and precuneus). No clinical seizures were observed during the course of the experiment. This study was approved by the SAHZU committee on Human Research.

### Experimental task

Each trial began with a video of animals, which lasted for 5-6 s. These videos were validated in previous studies (e.g., Zuo et al., 2020). Subsequently, at the memory test, the participants had to determine which image extracted from the video was presented earlier within a 10-second timeframe (temporal order judgment, TOJ). Following the memory test, a fixation cross was presented for 5 s. In the final phase, participants reported their confidence level on a scale ranging from 1 to 4. The number of trials completed by each participant varied, with a range of 120-240 trials (4-6 sessions, 30- 40 trials per session). Initiation of each trial was self-paced and the participants were allowed to take breaks between sessions. To exclude epileptic activity, we only included patients who did not experience seizures or pre-seizure activity within 24 hours.

### Intracranial recordings and electrode localization

Recording sites in depth electrodes implanted in the hippocampus and precuneus were 2 mm platinum cylinders with 1.5 mm intercontact spacing and a diameter of 0.8 mm (Fig.1b). Before electrode implantation, T1-weighted Magnetic Resonance Imaging (MRI) using a 3.0-T scanner and computed tomography were obtained for each patient. The locations of the electrodes were determined based on clinical standards, utilizing the SINOVATION software. We analyzed only electrodes localized to the hippocampus and precuneus in this study. Electrode localization was calculated using the Montreal Neurological Institute (MNI) coordinates, with the AAL3 atlas applied for further visualization. The number of electrodes for each patient is shown in the Supplementary table S1.

### Preprocessing

All data analyses were performed in MATLAB 2020a using EEGLAB, procedure from GitHub and Self-defining script. The sample rate of signal was at 2000 Hz. First, we extracted markers of each event, excluding trials with missing or wrong markers from further analysis. We selected electrodes from the same region based on MNI coordinates before bipolar referencing (either the hippocampus or precuneus, see Supplementary Table1). Then bipolar reference was applied to raw data of hippocampus and precuneus by measuring the difference between adjacent recording sites (i.e., 1.5 mm apart due to the electrode design). We used FIR band-stop filters to remove 50-Hz and 100-Hz power line interference. We focused on three events in the experimental task: 1. TOJ test (10s, retrieval); 2. Rest/Fixation (5s); 3. Report confidence level (4s, confidence rating).

### Data analysis

#### Generalized Linear Model (GLM)

To determine whether the neural signals are associated with metacognition, the time- series power (P) was regressed against several relevant factors (R). These factors included confidence level, confidence reaction time, response accuracy (correct/incorrect), retrieval reaction time, touch side (left/right), and video category:

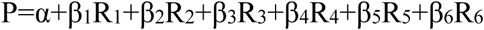

To assess the contribution of the different frequency bands to the confidence rating, confidence levels(R) were regressed separately across trials against normalized envelope of power P (Lopez-Persem et al., 2020) for each frequency band (delta: 1– 4Hz, theta: 4–8Hz, alpha: 8–14 Hz, beta: 14–35 Hz, low-gamma: 35-70 Hz, high- gamma: 70-150 Hz), averaged over time between 0 and 1.5 s before reporting confidence level:

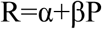

with α corresponding to the intercept. The significance of the regression estimates β was assessed across recording sites using two-sided, one-sample, Student’s t-tests.

To further determine whether other frequency bands provided additional information, we compared the following 7 GLMs:

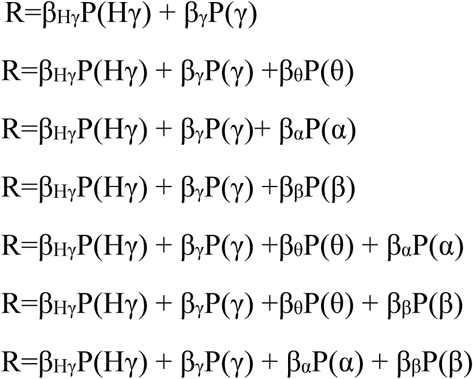

The model comparison was conducted using the Variational Bayesian Analysis (VBA) toolbox. Log-model evidence obtained in each recording site was taken to a group-level, random-effect, Bayesian model selection (RFX-BMS) procedure (Rigoux et al., 2014; Stephan et al., 2009). The RFX-BMS provided an exceedance probability (Xp) that measures the likelihood of a given model being more frequently implemented relative to all the others considered in the model space in the population from which samples are drawn.

#### Correlation Analysis

We used Pearson correlation to examine the relationship of high gamma activity between the hippocampus and precuneus. For correlation count, we firstly extracted the high-gamma (70-150Hz) band from both the hippocampus and precuneus. We then defined a window length of 0.02s, step by 0.01s (50% overlapping), and calculated the Pearson correlation coefficient for each window. To ensure the sensitivity and specificity of the results, we considered the signals to be relevant only when all 4 windows (equivalent to 0.04 s, at least 2 cycles of high gamma) met the minimum criteria of Pearson correlation (r > 0.8, p < 0.05). The occurrence of highly relevant signals during confidence rating was counted. We quantified the count of high correlation bands within the whole confidence rating stage. As a control, we shuffled the trials to obtain a baseline for each participant for statistical inference. The shuffling criteria were as follows: 1) random matching of signals from different trials, 2) a minimum interval of 10 trials between shuffled signals. Finally, we applied permutation test for origin data and shuffled data to detect significance (n=1000). As for the H-P correlation coefficient within time series, a window width of 0.02 s was also utilized to compute the Pearson correlation coefficient of high gamma frequency bands at each time window, averaging it according to the 0.1s window.

#### Measurement of metacognitive indices

We computed type 1 sensitivity (d’) and type 2 sensitivity (meta-d’), wherein meta-d’ indicates metacognitive sensitivity (the ability to discriminate between correct and incorrect judgments) and d’ represents memory performance according to signal detection theory (Green and Swets, 1966; Maniscalco and Lau, 2012). We divided within-subjects trials by their confidence levels into two categories for analysis: High confidence (3-4), Low confidence (1-2). In neural data, we computed an interaction index [(Neural data High confidence – Neural data Low confidence) - (Neural data Correct – Neural data Incorrect)] to reflect metacognitive process and an accuracy index (Neural data Correct – Neural data Incorrect) to reflect memory process.

#### Power Spectral Density (PSD)

During the confidence rating stages, we calculated the power spectral density (PSD) using the Welch periodogram method (pwelch function in MATLAB) for the 1 s interval preceding the confidence rating. A Hanning window of 0.1 s with 50% overlap was applied to minimize edge effects. Subsequently, we calculated the z-score for each electrode, followed by the computation of the accuracy index (Hγ _Correct_ – Hγ _Incorrect_) and interaction index [(Hγ _High confidence_ – Hγ _Low confidence_) - (Hγ _Correct_ – Hγ _Incorrect_)].

#### Time–frequency analysis

Time–frequency analyses were carried out using the Morlet wavelet transform method (cwt function in MATLAB) with default scale. We then computed the corresponding T values for each electrode.

## ACKNOWLEDGEMENT

We extend our gratitude to the patients for their cooperation, as well as to the medical staff involved in the clinical treatment and the technicians in the EEG laboratories for their valuable contributions.

## AUTHOR CONTRIBUTIONS

H.J.J and S.C.K. conceived the study and designed the experiment. K.P, X.X.G and H.Y.Y analyzed the data. J.M.Z supervised the surgeries and experiments. K.P and Z.L.Z. ran the experiments. Z.Z., J.M.Z, and H.J.J performed the surgeries and contributed to electrode localization. K.P and H.J.J wrote the paper. H.W, Y.C, D.S-P, and S.C.K further interpreted the data and contributed to the writing by reviewing and editing the manuscript.

## COMPETING INTERESTS

The authors declare no competing financial interests.

## FUNDING

S.C.K. received support from the National Natural Science Foundation of China (grant number: 32071060) and Jiangsu Provincial Department of Science and Technology (grant number: BK20221267).

## Notes

### Competing Interest Statement

The authors have declared no competing interest.

### Summary of Updates

Fig4 is in the wrong order

## REFERENCES

1. Allen, M., Glen, J. C., Müllensiefen, D., Schwarzkopf, D. S., Fardo, F., Frank, D., Callaghan, M. F., & Rees, G. (2017). Metacognitive ability correlates with hippocampal and prefrontal microstructure. NeuroImage, 149, 415–423. 10.1016/j.neuroimage.2017.02.008

2. Baird, B., Smallwood, J., Gorgolewski, K. J., & Margulies, D. S. (2013). Medial and lateral networks in anterior prefrontal cortex support metacognitive ability for memory and perception. Journal of Neuroscience, 33(42), 16657–16665. 10.1523/JNEUROSCI.0786-13.2013

3. Bicanski, A., & Burgess, N. (2018). A neural-level model of spatial memory and imagery. ELife, 7. 10.7554/eLife.33752

4. Boldt, A., & Gilbert, S. J. (2022). Partially Overlapping Neural Correlates of Metacognitive Monitoring and Metacognitive Control. Journal of Neuroscience, 42(17), 3622–3635. 10.1523/JNEUROSCI.1326-21.2022

5. Burke, J. F., Long, N. M., Zaghloul, K. A., Sharan, A. D., Sperling, M. R., & Kahana, M. J. (2014). NeuroImage Human intracranial high-frequency activity maps episodic memory formation in space and time. NeuroImage, 85, 834–843. 10.1016/j.neuroimage.2013.06.067

6. Cai, Y., Jin, Z., Zhai, C., Wang, H., Wang, J., Tang, Y., & Kwok, S. C. (2022). Time-sensitive prefrontal involvement in associating confidence with task performance illustrates metacognitive introspection in monkeys. Communications Biology, 5(1). 10.1038/s42003-022-03762-6

7. Das, A., & Menon, V. (2023). Hippocampal-parietal cortex causal information flow in human episodic memory : replication across three experiments.

8. Desender, K., Ridderinkhof, K. R., & Murphy, P. R. (2021). Understanding neural signals of post-decisional performance monitoring: An integrative review. ELife, 10, 1–21. 10.7554/eLife.67556

9. Flavell, J. H. (1979). Metacognition and cognitive monitoring: A new area of cognitive– developmental inquiry. American Psychologist, 34(10), 906–911.

10. Fleming, Stephen M; Weil, Rimona S; Nagy, Zoltan; Dolan, Raymond J; & Rees, G. (2010). Tackling Attack Detection and Incident Response. Science, 329(April), 1541–1543. 10.1126/science.1191883.Relating

11. Fleming, S. M. (2024). Metacognition and Confidence: A Review and Synthesis. Annual Review of Psychology, 75(1), 1–28. 10.1146/annurev-psych-022423-032425

12. Foudil, S. A., Kwok, S. C., & Macaluso, E. (2020). Context-dependent coding of temporal distance between cinematic events in the human precuneus. Journal of Neuroscience, 40(10). 10.1523/JNEUROSCI.2296-19.2020

13. Futing Zou, Sze Chai Kwok; Distinct Generation of Subjective Vividness and Confidence during Naturalistic Memory Retrieval in Angular Gyrus. J Cogn Neurosci 2022; 34 (6): 988–1000. doi: 10.1162/jocn_a_01838

14. Green, D.M., Swets, J.A., 1966. Signal Detection Theory and Psychophysics. John Wiley, Oxford.

15. Haque, R. U., Inati, S. K., Levey, A. I., & Zaghloul, K. A. (n.d.). episodic memory retrieval. 2020, 1–14. 10.1038/s41467-020-19828-0

16. Hebart, M. N., Schriever, Y., Donner, T. H., & Haynes, J. D. (2016). The Relationship between Perceptual Decision Variables and Confidence in the Human Brain. Cerebral Cortex, 26(1), 118–130. 10.1093/cercor/bhu181

17. Hebscher, M., Ibrahim, C., & Gilboa, A. (2020). Precuneus stimulation alters the neural dynamics of autobiographical memory retrieval. NeuroImage, 210(June 2019), 116575. 10.1016/j.neuroimage.2020.116575

18. Hebscher, M., Meltzer, J. A., & Gilboa, A. (2019). A causal role for the precuneus in network-wide theta and gamma oscillatory activity during complex memory retrieval. ELife, 8, 1–20. 10.7554/eLife.43114

19. Heereman, J., Walter, H., & Heekeren, H. R. (2015). A task-independent neural representation of subjective certainty in visual perception. Frontiers in Human Neuroscience, 9(OCT). 10.3389/fnhum.2015.00551

20. Koch G, Casula EP, Bonnì S, Borghi I, Assogna M, Minei M, Pellicciari MC, Motta C, D’Acunto A, Porrazzini F, Maiella M, Ferrari C, Caltagirone C, Santarnecchi E, Bozzali M, Martorana A. Precuneus magnetic stimulation for Alzheimer’s disease: a randomized, sham-controlled trial. Brain. 2022 Nov 21;145(11):3776–3786. doi: 10.1093/brain/awac285. PMID: 36281767; PMCID: PMC9679166.

21. Kwok, S. C., & Buckley, M. J. (2010). Fornix transection selectively impairs fast learning of conditional visuospatial discriminations. Hippocampus, 20(3), 413–422. 10.1002/hipo.20643

22. Kwok, S. C., Cai, Y., & Buckley, M. J. (2019). Mnemonic introspection in macaques is dependent on superior dorsolateral prefrontal cortex but not orbitofrontal cortex. Journal of Neuroscience, 39(30), 5922–5934. 10.1523/JNEUROSCI.0330-19.2019

23. Kwok, S. C., Mitchell, A. S., & Buckley, M. J. (2015). Adaptability to changes in temporal structure is fornix-dependent. Learning and Memory, 22(8), 354–359. 10.1101/lm.038851.115

24. Lopez-Persem, A., Bastin, J., Petton, M., Abitbol, R., Lehongre, K., Adam, C., Navarro, V., Rheims, S., Kahane, P., Domenech, P., & Pessiglione, M. (2020). Four core properties of the human brain valuation system demonstrated in intracranial signals. Nature Neuroscience, 23(5), 664 – 675. 10.1038/s41593-020-0615-9

25. Maniscalco, B., & Lau, H. (2012). A signal detection theoretic approach for estimating metacognitive sensitivity from confidence ratings. Consciousness and Cognition, 21(1), 422–430. 10.1016/j.concog.2011.09.021

26. Manippa, V., Palmisano, A., Nitsche, M. A., Filardi, M., Vilella, D., Logroscino, G., & Rivolta, D. (2024). Cognitive and Neuropathophysiological Outcomes of Gamma-tACS in Dementia: A Systematic Review. Neuropsychology Review, 34(1), 338–361. 10.1007/s11065-023-09589-0

27. Mart, C. (2021). The hippocampus as the switchboard between perception and memory. 118(50). 10.1073/pnas.2114171118/-/DCSupplemental.Published

28. McCurdy, L. Y., Maniscalco, B., Metcalfe, J., Liu, K. Y., de Lange, F. P., & Lau, H. (2013). Anatomical coupling between distinct metacognitive systems for memory and visual perception. Journal of Neuroscience, 33(5), 1897–1906. 10.1523/JNEUROSCI.1890-12.2013

29. Miyamoto K, Osada T, Setsuie R, Takeda M, Tamura K, Adachi Y, Miyashita Y. Causal neural network of metamemory for retrospection in primates. Science. 2017 Jan 13;355(6321):188-193. doi: 10.1126/science.aal0162. PMID: 28082592.

30. Miyamoto, K., Setsuie, R., Osada, T., & Miyashita, Y. (2018). Reversible Silencing of the Frontopolar Cortex Selectively Impairs Metacognitive Judgment on Non-experience in Primates. Neuron, 97(4), 980–989.e6. 10.1016/j.neuron.2017.12.040

31. Miyoshi, K., Webb, T., Rahnev, D., & Lau, H. (2024). Confidence and metacognition. In Reference Module in Neuroscience and Biobehavioral Psychology. Elsevier. 10.1016/b978-0-12-820480-1.00049-8

32. Mograbi DC, Ferri CP, Sosa AL, Stewart R, Laks J, Brown R, Morris RG. Unawareness of memory impairment in dementia: a population-based study. Int Psychogeriatr. 2012 Jun;24(6):931–9. doi: 10.1017/S1041610211002730. Epub 2012 Jan 17. PMID: 22251835.

33. Morales, J., Lau, H., & Fleming, S. M. (2018). Domain-general and domain-specific patterns of activity supporting metacognition in human prefrontal cortex. Journal of Neuroscience, 38(14), 3534–3546. 10.1523/JNEUROSCI.2360-17.2018

34. Morris, Robin G, and Kristin Hannesdottir, ’Loss of ‘Awareness’ in Alzheimer’s Disease’, i n Robin G Morris, and James T Becker (eds), Cognitive Neuropsychology of Alzheimer’s Disease (Oxford, 2004; online edn, Oxford Academic, 31 Oct. 2023), 10.1093/o so/9780198508304.003.0017, accessed 18 Aug. 2024.

35. Pereira, M., Megevand, P., Tan, M. X., Chang, W., Wang, S., Rezai, A., Seeck, M., Corniola, M., Momjian, S., Bernasconi, F., Blanke, O., & Faivre, N. (2021). Evidence accumulation relates to perceptual consciousness and monitoring. Nature Communications, 12(1), 1–11. 10.1038/s41467-021-23540-y

36. Pudhiyidath, A., Morton, N. W., Duran, R. V., Schapiro, A. C., Momennejad, I., Hinojosa-Rowland, D. M., Molitor, R. J., & Preston, A. R. (2022). Representations of Temporal Community Structure in Hippocampus and Precuneus Predict Inductive Reasoning Decisions. Journal of Cognitive Neuroscience, 34(10), 1736–1760. 10.1162/jocn_a_01864

37. Ren, Y., Nguyen, V. T., Sonkusare, S., Lv, J., Pang, T., Guo, L., Eickhoff, S. B., Breakspear, M., & Guo, C. C. (2018). Effective connectivity of the anterior hippocampus predicts recollection confidence during natural memory retrieval. Nature Communications, 9(1). 10.1038/s41467-018-07325-4

38. Rigoux, L., Stephan, K. E., Friston, K. J., & Daunizeau, J. (2014). Bayesian model selection for group studies - Revisited. NeuroImage, 84, 971 – 985. 10.1016/j.neuroimage.2013.08.065

39. Richter, F. R., Cooper, R. A., Bays, P. M., & Simons, J. S. (2016). Distinct neural mechanisms underlie the success, precision, and vividness of episodic memory. ELife, 5(OCTOBER2016), 1–18. 10.7554/eLife.18260

40. Rutishauser, U., Ye, S., Koroma, M., Tudusciuc, O., Ross, I. B., Chung, J. M., & Mamelak, A. N. (2015). Representation of retrieval confidence by single neurons in the human medial temporal lobe. Nature Neuroscience, 18(7), 1041–1050. 10.1038/nn.4041

41. Shekhar, M., & Rahnev, D. (2018). Distinguishing the roles of dorsolateral and anterior PFC in visual metacognition. Journal of Neuroscience, 38(22), 5078–5087. 10.1523/JNEUROSCI.3484-17.2018

42. Staresina, B. P., Michelmann, S., Bonnefond, M., Jensen, O., Axmacher, N., & Fell, J. (2016). Hippocampal pattern completion is linked to gamma power increases and alpha power decreases during recollection. 1–18. 10.7554/eLife.17397

43. Stephan, K. E., Penny, W. D., Daunizeau, J., Moran, R. J., & Friston, K. J. (2009). Bayesian model selection for group studies. NeuroImage, 46(4), 1004 – 1017. 10.1016/j.neuroimage.2009.03.025

44. Tanglay, O., Young, I. M., Dadario, N. B., Briggs, R. G., Fonseka, R. D., Dhanaraj, V., Hormovas, J., Lin, Y. H., & Sughrue, M. E. (2022). Anatomy and white-matter connections of the precuneus. In Brain Imaging and Behavior (Vol. 16, Issue 2, pp. 574–586). 10.1007/s11682-021-00529-1

45. Voytek B, D’Esposito M, Crone N, Knight RT. A method for event-related phase/amplitude coupling. Neuroimage. 2013;64:416–424. doi:10.1016/j.neuroimage.2012.09.023

46. Xue, K., Zheng, Y., Rafiei, F., & Rahnev, D. (2023). The timing of confidence computations in human prefrontal cortex. Cortex, 1–28. 10.1016/j.cortex.2023.08.009

47. Ye, Q., Zou, F., Dayan, M., Lau, H., Hu, Y., & Kwok, S. C. (2019). Individual susceptibility to TMS affirms the precuneal role in meta-memory upon recollection. Brain Structure and Function, 224(7), 2407–2419. 10.1007/s00429-019-01909-6

48. Ye, Q., Zou, F., Lau, H., Hu, Y., & Kwok, S. C. (2018). Causal evidence for mnemonic metacognition in human precuneus. Journal of Neuroscience, 38(28), 6379–6387. 10.1523/JNEUROSCI.0660-18.2018

49. Zheng, Y., Bo, B., Wang, D., Liu, Y., Gilbert, S. J., Li, Y., & Kwok, S. C. (2024). Fornix and Uncinate Fasciculus Support Metacognition-Driven Cognitive Offloading. 21, 1–26.

50. Zheng, Y., Wang, D., Ye, Q., Zou, F., Li, Y., & Kwok, S. C. (2021). Diffusion property and functional connectivity of superior longitudinal fasciculus underpin human metacognition. Neuropsychologia,156 (October 2020), 107847. 10.1016/j.neuropsychologia.2021.107847

51. Zheng, A., Montez, D. F., Marek, S., Gilmore, A. W., Newbold, D. J., Laumann, T. O., Kay, B. P., Seider, N. A., Van, A. N., Hampton, J. M., Alexopoulos, D., Schlaggar, B. L., Sylvester, C. M., Greene, D. J., Shimony, J. S., Nelson, S. M., Wig, G. S., Gratton, C., McDermott, K. B., … Dosenbach, N. U. F. (2021). Parallel hippocampal-parietal circuits for self- And goal-oriented processing. Proceedings of the National Academy of Sciences of the United States of America, 118(34). 10.1073/pnas.2101743118

52. Zou, Futing, and Sze Chai Kwok. “Distinct Generation of Subjective Vividness and Confidence during Naturalistic Memory Retrieval in Angular Gyrus.” Journal of cognitive neuroscience vol. 34,6 (2022): 988–1000. doi:10.1162/jocn_a_01838

53. Zuo, S., Wang, L., Shin, J. H., Cai, Y., Lee, S. W., Appiah, K., Zhou, Y. Di, & Kwok, S. C. (2020). Behavioral evidence for memory replay of video episodes in the macaque. ELife, 9, 1–23. 10.7554/eLife.54519

