## Supplemental Figure.S1;Supplemental Figure.S2;Supplemental Figure.S3;Supplemental Fig.S4;Supplemental Table 1 for "Components of Mnemonic Metacognition Constitutionally Supported by High Gamma Activity Between the Precuneus and Hippocampus" (from another version of this preprint: version 2, 2024-10-20)

Correspondence

*Table of Content:*

1.[Supplementary Figure	3](#_Toc177133435)

1.1. [Figure. S1. The correlation between high gamma activity in the precuneus and hippocampus with both d' and meta-d'.	3](#_Toc177133436)

1.2. [Figure. S2. The probability density comparing shuffled and original data of 10 subject.	4](#_Toc177133437)

1.3. [Figure. S3. The relationship between correlation count of high gamma activity in the hippocampus and precuneus with both d' and meta-d'.	5](#_Toc177133438)

1.4. [Figure. S4. Phase amplitude coupling (PAC) and mean modulation index in three stages.](#_Toc177133438)	6

2.[Supplementary Table](#_Toc177133439)	7

2.1. [Table. S1. Basic information and electrodes of patients.](#_Toc177133440)	7

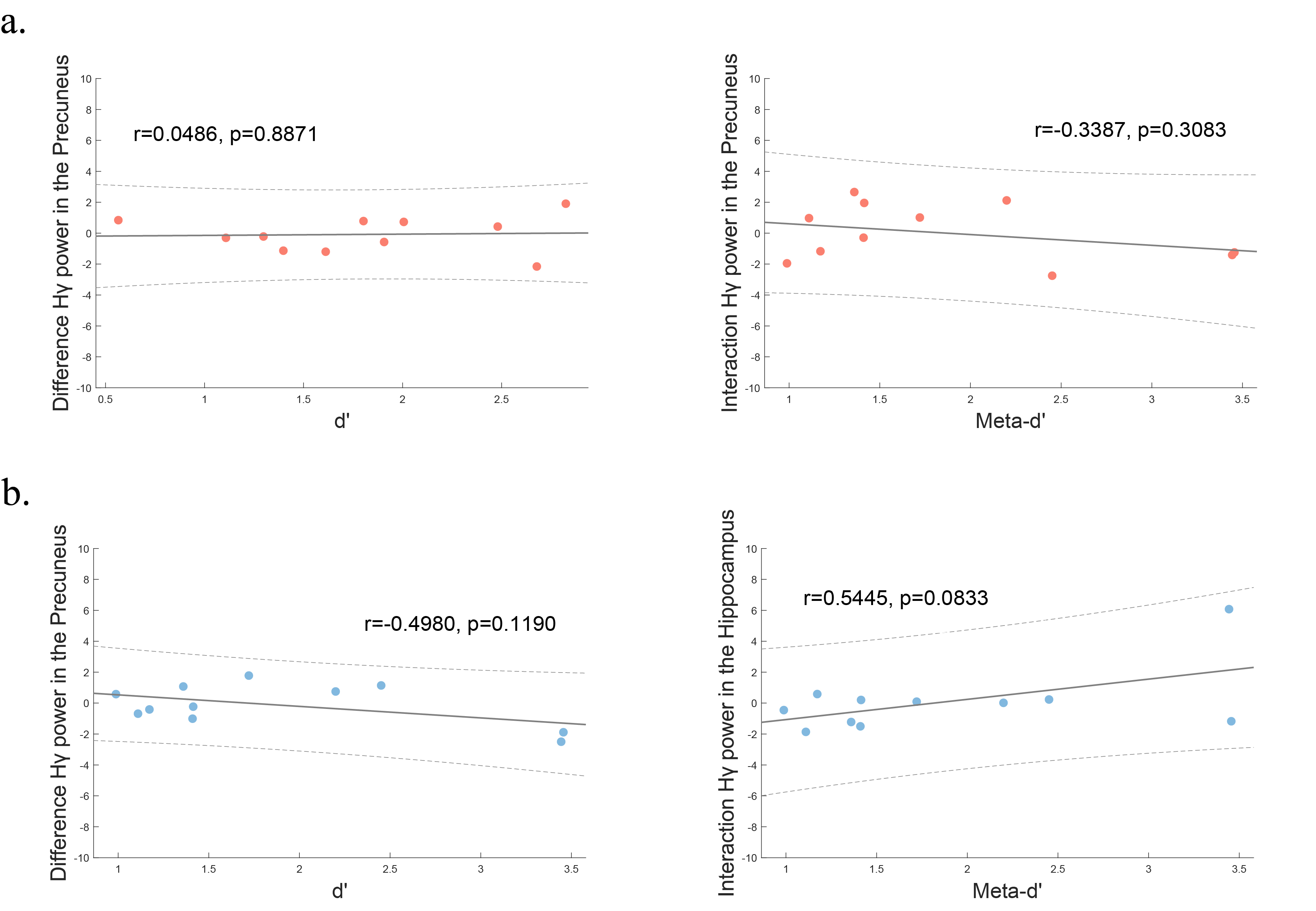

**Figure. S1. The correlation between high gamma activity in the precuneus and hippocampus with both d' and meta-d'. (a)** The left panel illustrates the correlation around -0.8~-0.5s time window between the neural accuracy index (Hγ _Correct_ – Hγ _Incorrect_) in the precuneus and behavioral type 1 sensitivity (d') (Pearson correlation, r=0.0486, p=0.8871) within the hippocampus's significant time window (i.e., -0.8~-0.5s), whereas the right panel depicts the correlation between the interaction index [(Hγ _High confidence_ – Hγ _Low confidence_) - (Hγ _Correct_ – Hγ _Incorrect_)] and the type 2 sensitivity (meta-d') (Pearson correlation, r = -0.3387, p = 0.3083) within the same time window. **(b)** The left panel illustrates the correlation around -0.3~0s time window between the neural accuracy index (Hγ _Correct_ – Hγ _Incorrect_) in the hippocampus and behavioral type 1 sensitivity (d') (Pearson correlation, r = 0.0486, p = 0.8871) within the precuneus’ significant time window (i.e., -0.3~0s), whereas the right panel depicts the correlation between the neural interaction index [(Hγ _High confidence_ – Hγ _Low confidence_) - (Hγ _Correct_ – Hγ _Incorrect_)] and the behavioral type 2 sensitivity (meta-d') (Pearson correlation, r = -0.3387, p = 0.3083) within the same time window.

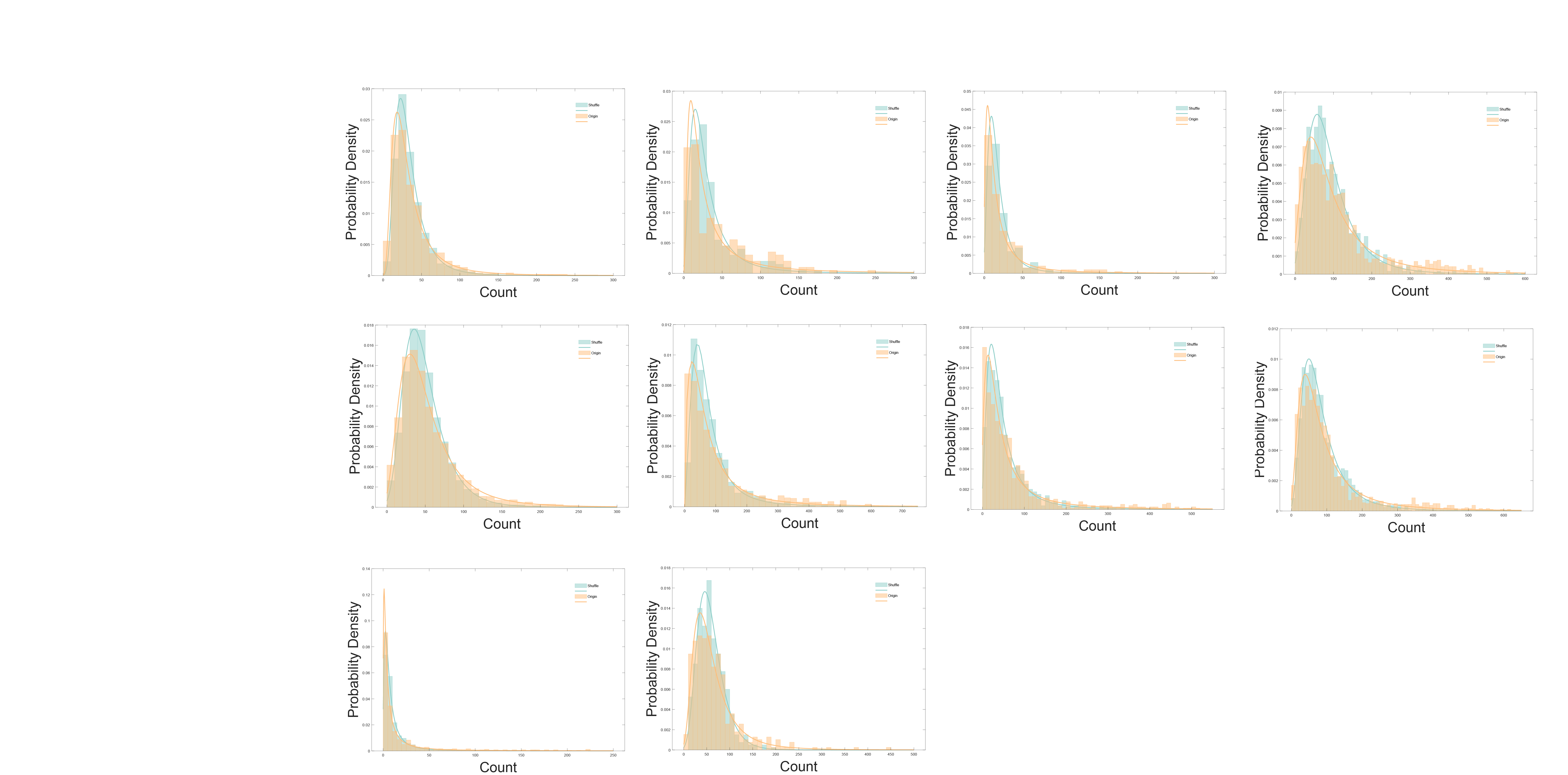

**Figure. S2. The probability density comparing shuffled and original data of 10 participants (in addition to the participant shown in Figure 5B).** All **participants** demonstrated significant differences in the original data when compared to the shuffled data (permutation test, n=1000).

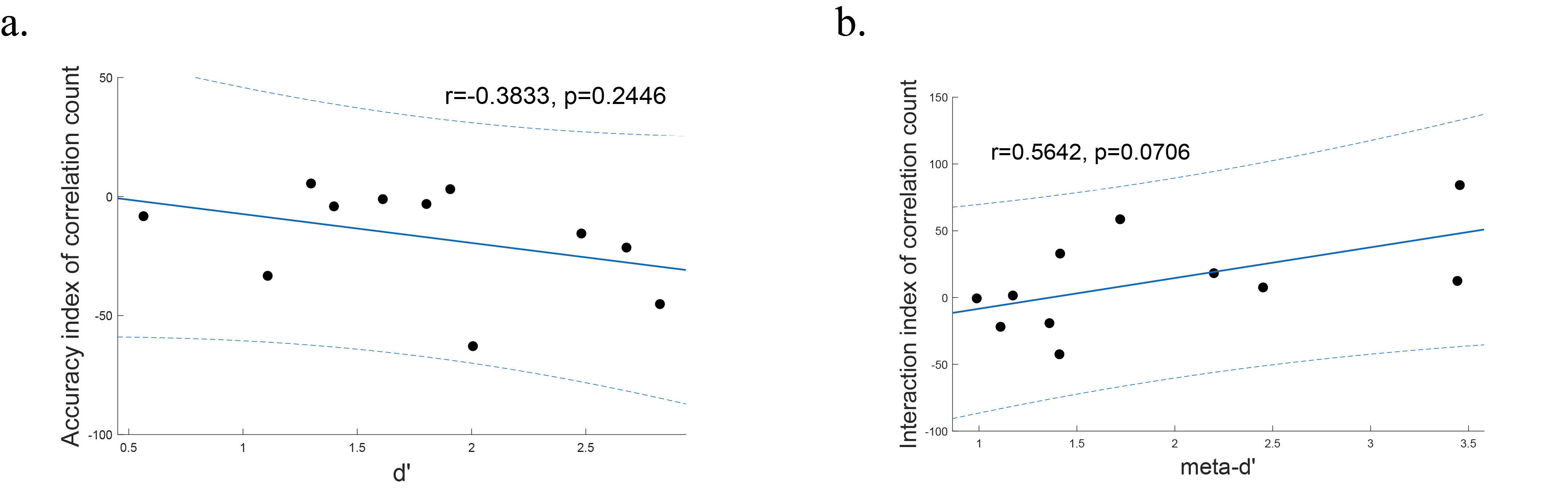

**Figure. S3. The relationship between correlation count of high gamma activity in the hippocampus and precuneus with d' and meta-d'. (a)** The correlation between neural accuracy index (Count _Correct_ – Count _Incorrect_) of correlation count and behavioral type 1 sensitivity (d') (Pearson correlation, r = -0.3833, p = 0.2446). (b) The correlation between neural interaction index [(Count _High confidence_ – Count _Low confidence_) - (Count _Correct_ – Count _Incorrect_)] of correlation count and behavioral type 2 sensitivity (meta-d') (Pearson correlation, r = 0.5642, p = 0.0706).

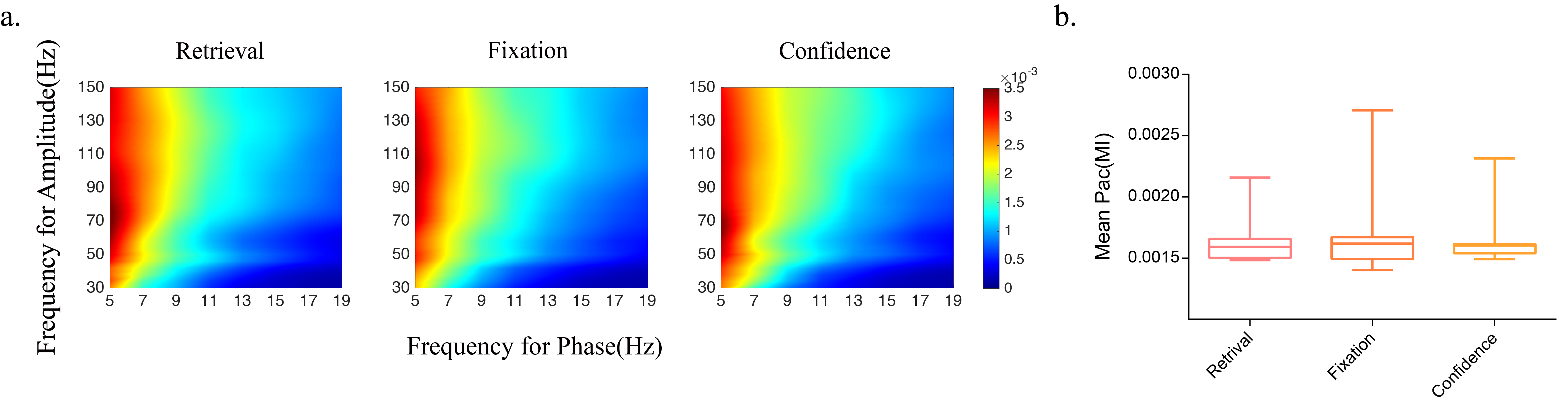

**Figure. S4. Phase amplitude coupling (PAC) and mean modulation index in three stages.** (a) Representative example of PAC observed in precuneus (phase) and hippocampus (amplitude) of one participant in retrieval (left), fixation (middle), confidence (right). The warmest colors represent the strongest coupling. The amplitude frequency ranges from 30-150Hz, and phase frequency ranges from 5-19Hz. (b) Box plot shows the mean modulation index in the three periods (Friedman test, p > 0.05).

**Table. S1. Basic demongraphy and electrode information of patients.**

| Patient# | Age | Sex | #Electrocode | #Total recording  sites | Cerebral hemisphere | #Selected sites of hippocampus | #Selected sites of precuneus |
| --- | --- | --- | --- | --- | --- | --- | --- |
| 1 | 12 | M | 12 | 145 | Left | 3 | 3 |
| 2 | 28 | M | 12 | 148 | Right | 4 | 4 |
| 3 | 17 | M | 10 | 118 | Left | 3 | 4 |
| 4 | 20 | M | 11 | 112 | Left | 2 | 2 |
| 5 | 26 | F | 11 | 120 | Right | 3 | 4 |
| 6 | 16 | M | 14 | 158 | Left | 3 | 5 |
| 7 | 39 | M | 12 | 148 | Left | 7 | 6 |
| 8 | 37 | M | 14 | 172 | Right | 4 | 5 |
| 9 | 34 | M | 15 | 202 | Left | 2 | 3 |
| 10 | 14 | F | 12 | 134 | Left | 2 | 2 |
| 11 | 31 | F | 11 | 144 | Left | 2 | 4 |
